# Qualification of Identity, Quality-Attribute Monitoring and New Peak Detection in An Improved Multi-Attribute Method with Lys-C Digestion for Characterization and Quality Control of Therapeutic Monoclonal Antibodies

**DOI:** 10.1101/2022.05.01.490194

**Authors:** Xuanwen Li, Nicholas A. Pierson, Xiaoqing Hua, Bhumit A. Patel, Michael H. Olma, Christopher A. Strulson, Simon Letarte, Douglas D. Richardson

## Abstract

The use of Multi-attribute method (MAM) for identity and purity testing of biopharmaceuticals offers the ability to complement and replace multiple conventional analytical technologies with a single mass spectrometry (MS) method. Method qualification and phase-appropriate validation is one major consideration for the implementation of MAM in a current Good Manufacturing Practice (cGMP) environment. We developed an improved MAM workflow with optimized sample preparation using Lys-C digestion for therapeutic monoclonal antibodies. In this study, we qualified the enhanced MAM workflow for mAb-1 identity, product quality attributes (PQAs) monitoring and new peak detection (NPD). The qualification results demonstrated the full potential of the MAM for its intended use in mAb-1 characterization and quality control in regulated labs. To the best of our knowledge, this is the first report of MAM qualification for mAb identity, PQA monitoring, and new peak detection (NPD) in a single assay, featuring 1) the first full qualification of MAM using Lys-C digestion without desalting using a high-resolution MS, 2) a new approach for mAb identity testing using MAM, and 3) the first qualification of NPD for MAM. The developed MAM workflow and the approaches for MAM qualification may serve as a reference for other labs in the industry.

## Introduction

The pharmaceutical industry is transitioning toward Quality by Design (QbD) for the manufacturing of biotherapeutics. QbD is also the expectation from heath authorities, where quality of biotherapeutics must be built into the process used to produce and test therapeutics.^1^ Analytical method lifecycle management typically begins with an analytical target profile (ATP) followed by method development, phase-appropriate qualification/validation, method transfer, and operational use in routine testing.^1–2^ Optimization occurs throughout the method’s lifecycle starting in development and moving to method robustness, once there is a better understanding of experimental parameters. One aspect of QbD is constant method improvement which is a critical aspect of the analytical method life cycle management to meet additional quality needs, such as analytical support of co-formulated drug products.

The current state of releasing biotherapeutics from a quality control (QC) lab involves a large variety of conventional assays,^3–5^ including capillary electrophoresis (CE) or high performance liquid chromatography (HPLC) based purity assays, as well as host cell proteins (HCPs) or protein A based enzyme-linked immunosorbent assay (ELISA) for process-related impurities. If one or more of these assays fails to meet its release/stability specifications, orthogonal attribute assays like MS-based assays can be employed to assist the root-cause investigation. Building from commercial QC success with MS for other modalities (e.g., small molecules and peptides) provides a key opportunity for its use in large molecule characterization.^6–7^ Application of MS assays for monoclonal antibodies, including intact/reduced/subunit mass, and non-reduced/reduced/focused peptide mapping as primary methods for in-process, release, and stability in clinical manufacturing is desired to achieve enhanced product understanding along with increased efficiency.^1, 6–10^

Reduced peptide mapping based multi-attribute method (MAM) is a MS-based method that can be utilized in a QC lab as an identity test as well as a purity test, to monitor known CQAs and to detect potential impurities by new peak detection (NPD).^6^ A successful transition of MAM as a primary assay into a GMP lab requires a comprehensive characterization of the therapeutic protein for risk assessment, method validation, comparison to conventional assays, and NPD.^6^ With extensive characterization of biological molecules and an understanding of the relationship between global structure changes and specific post-translational modifications (PTM), MAM is a completely orthogonal assay to some conventional techniques and has the potential to be a primary method for release and stability testing in QC labs for biotherapeutics.^1, 6–7, 10–20^

Co-formulated therapies are designed to effectively enhance clinical benefit with reduced medication errors and enhanced convenience for patients, as well as extend intellectual property rights for pharmaceutical companies.^21^ Co-formulation also increases the complexity of drug product development and creates additional challenges for analytical characterization and quality control of drug products, especially for co-formulated antibodies with similar physicochemical properties and widely disparate concentrations (e.g., 20:1 ratio).^22–23^ For example, many traditional identity (ID) assays, including ion exchange chromatography (IEX), potency-based ELISA, and ultraviolet (UV) based peptide mapping methods have their unique challenges to confidently identify co-formulated mAbs. The application of MAM for ID of co-formulated mAbs seems straightforward, but has not been reported.

In previous work, we developed a reduced peptide mapping based MAM with improved sample preparation using Lys-C digestion without desalting for mAbs and their co-formulations.^24^ We also optimized peptide separation for hydrophilic peptides, especially for a stability-indicating peptide, VSNK.^24^ In this study, we qualified the improved MAM assay for ID, PQA monitoring, and NPD in method development labs based on the guidance for validation of analytical procedures from International Conference on Harmonization (ICH) guideline Q2 (R1) as shown in Figure 1.^6^

**Figure 1:**
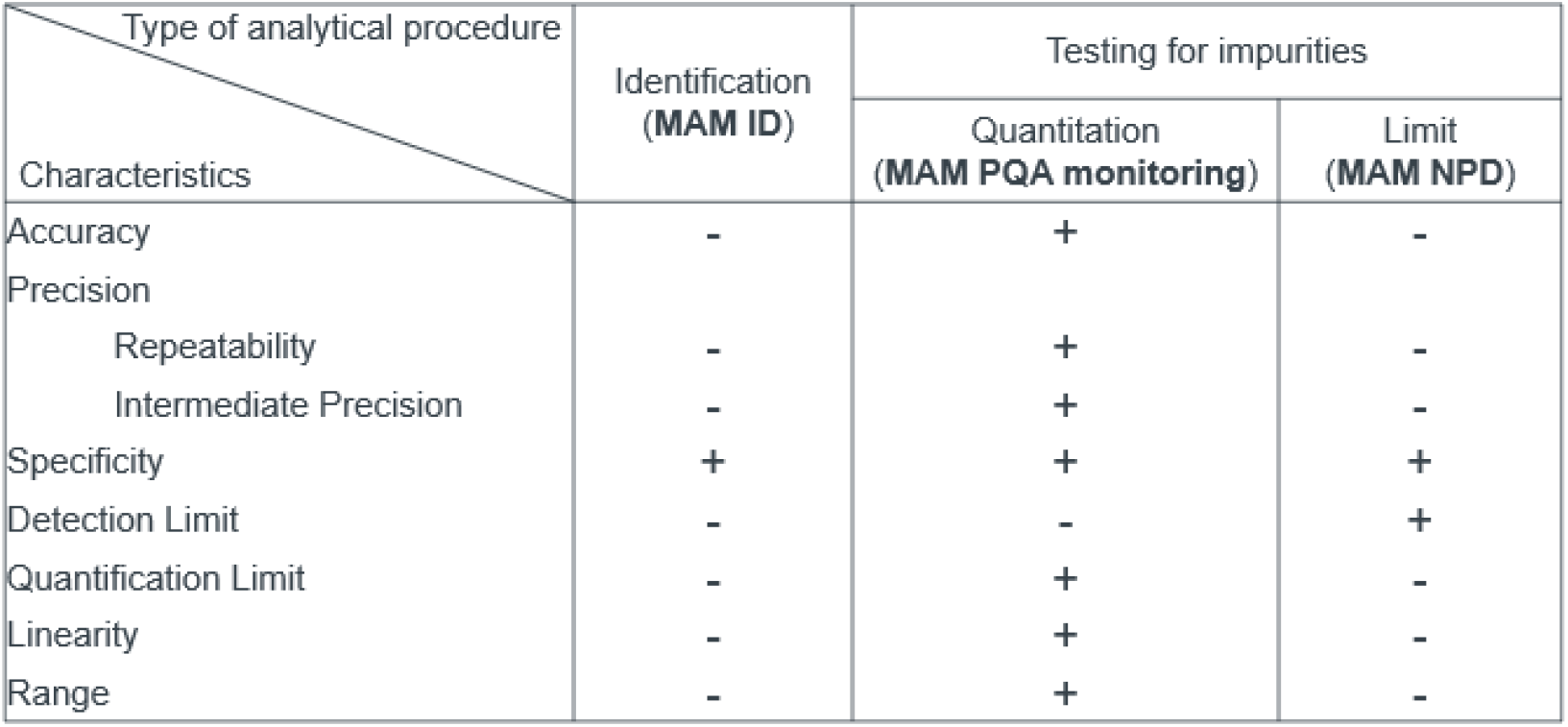
The MAM qualification strategy modified from ICH guideline Q2 (R1). MAM application for ID were qualified as an identification test. MAM application for PQA monitoring was qualified as a quantitation testing for impurities, and MAM application for NPD was qualified as a limit testing for impurities.

## Materials and Methods

### Materials

Dithiothreitol (DTT) and iodoacetamide (IAM), both in no-weigh formats, and Pierce™ peptide retention time calibration mixture (PRTC) were purchased from Thermo Pierce (Rockford, IL). Tris-HCl buffer (1 M, pH 8.0), guanidine-HCl (8 M), water (Optima™), acetonitrile (ACN, Optima™), trifluoracetic acid (TFA) were purchased from Thermo Fisher Scientific (Waltham, MA). Ethylenediaminetetra-acetic acid (EDTA, 0.5 M, pH 8.0) were purchased from Promega (Madison, WI). Lysyl endopeptidase (Lys-C) was purchased from Wako (Osaka, Japan). mAb-1 to mAb-7 are monoclonal antibodies in development at Merck & Co., Inc., Kenilworth, NJ, USA.

### Sample preparation

The protein digestions were conducted as previously reported.^24^ Briefly, a total of 20 μL of the diluted sample (containing 100 μg) was denatured and reduced in final solution (100 μL) containing approximately 6 M Guanidine-HCl, 50 mM Tris-HCl, 5 mM EDTA and 20 mM DTT. The samples were incubated in a thermomixer at 37°C with 300 rpm shaking for 30 min, then were alkylated with 5 μL of IAM (1 M) and protected from light at 25 °C for 30 min. After the block of unreacted IAM with 5 μL of DTT (200 mM), Lys-C enzyme (1:10 (wt: wt)) in 500 μL of 50 mM Tris was added to the protein sample. The samples were incubated in a thermomixer at 37°C for 60 min, and subsequently quenched with 15 μL of 20% TFA. The digested samples were analyzed by LC-MS within 24 h after sample digestion. Otherwise, digested samples were stored at −80 °C for future analysis.

### UPLC-MS based MAM and data analysis

The LC-MS was run as previously reported.^24^ The LC was run utilized a Waters Acquity UPLC H-Class Bio system with 20 μL (3.2 μg) of sample per injection onto a Waters HSS T3 column (100Å, 1.8 μm, 2.1 mm X 150 mm) with column temperature (40 ± 3 °C). Autosampler was set at 5°C. Mobile phase (MP)-A was 0.02% TFA in water and MP-B was 0.02% TFA in acetonitrile. The flow rate was 0.3 ml/min. The LC gradient started with 0.1% MP-B from 0 to 5 min, 0.1%-12% MP-B from 5 to 7 min, followed by an increase to 40% MP-B over the next 38 min. The column was washed with 98% MP-B for 4 min before returning to 0.1% MP-B for balancing. The run time of the entire gradient was 1 hour.

The MS detection was acquired with a Q Exactive™ or Exactive Plus™ Orbitrap MS (Thermo) using Chromeleon (7.2.10, Thermo). The MS parameters included spray voltage at 3.8 kV, capillary temperature at 250 °C, sheath gas at 35 (arbitrary units), aux gas at 10 (arbitrary units), S-Lens RF level at 50, aux gas heater temperature at 250 °C, scan range from 300 to 1800 m/z, AGC target at 1E6, maximum inject time at 200 ms, MS resolution at 140,000, mass acquisition time from 2.2 to 45.0 min, and lock mass with 391.2843 m/z.

For PQA quantification in Chromeleon (Thermo, Waltham, MA), the two or three most abundant charge states (each with the top two or three abundant isotope ions, only +1 charge state was used for the VSNK peptide) of a peptide (determined from development studies) were selected for data process and reporting. The pass score for composite scoring had the following 3 criteria: 1) isotopic dot product ≥ 0.8; 2) mass accuracy ≤ 5 ppm; and 3) peak apex alignment ≤ 0.5 min. The extracted ion chromatogram (EIC) of non-modified and modified peptides of a specific PQA were used to determine the percent of PTM using the formula [peak area of the extracted ion chromatogram (EIC) of the modified peptide] / [peak area of EIC of the modified peptide + peak area of EIC of the unmodified peptide] x 100.

The settings for NPD in Chromeleon included m/z min at 300, m/z max at 1800, m/z width at 5 ppm, RT start at 2.2 min, RT stop at 45 min, frame time width at 1 min, and Max number of frames at 5,000. The peak intensity thresholds at 0.01%, 0.1% and 0.3% of TIC were evaluated. The frame rules for NPD in Chromeleon included charge>1, PR element=0, PR size >1, and ratio is not between 0.2 and 5.0.

### Qualification for mAb-1 identity

The Peak Area Ratios (PARs) for Lys-C digested N-terminals of both heavy chain and light chain of mAb-1, −2, −3 and −4 were calculated by comparing them to the non-modified PENNYK peptide in the same run with the following formula:

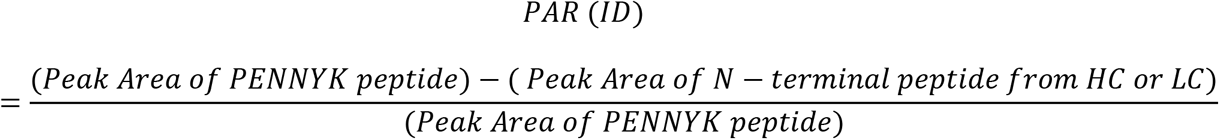

For positive identification of one or two mAb (as in co-formulations), the PARs of both the heavy-chain and light-chain peptides must be ≤ 0.97.

### Qualification for PQA monitoring

Study design and performance characteristics for the qualification of PQA monitoring were based on its intended use and phase of development.^6^ Pre-defined acceptance criteria of this qualification exercise were not set in the method development labs. For the MAM development of mAb-1, a risk assessment was performed.^6^ In the end, one methionine (M) oxidation site, two asparagine deamidation sites (N1 and N2), and the N-terminal and C-terminal variances of heavy chains were included in the Chromeleon workbook.

PQA monitoring was qualified for the characteristics of quantitation testing for impurities according to ICH Q2 (R1) guideline, which included specificity, linearity, range, quantitation limit, precision (repeatability and intermediate precision) and accuracy (Figure 1).

For specificity, a water injection, mAb-1 formulation and in-process buffers, mAb-1 reference standard (RS) and mAb-1 light-stressed sample were analyzed by MAM. The specificity of the method was evaluated by comparing the chromatograms of mAb-1 samples to other injections.

To evaluate repeatability, the mAb-1 RS was digested in 6 preparations and analyzed as one precision run. To evaluate intermediate precision, the precision run was performed with a second analyst on different days with the same or different instrument and same or different analytical column in two labs. A total of 6 precision runs were performed following the analyzing scheme as shown in the supplementary Table 1.

To test the linearity of the method, a range of volumes of mAb-1 digest (2, 5, 10, 20, 30, 40, and 50 μL) were injected into the LC-MS system. The injection volume varies between 10% and 250% of the target injection volume (20 μL). The accuracy of MAM for PQA quantification was determined by recovery, which was estimated using the detected peak area compared to the calculated peak area from the 100% assay level (20 μL) based on the injection volume. Assay range and limit of quantification (LOQ) were inferred from the linearity study.

The lower LOQ for each PQA was calculated according to the following formula:

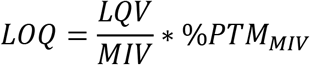

LQV (lowest quantification volume) was set as the lowest injection volume in range with acceptable linearity (R^2^≥0.99), accuracy (from 75% to 125% of calculated level) and precision (≤15% RSD). MIV (middle injection volume) was the standard injection volume (20 μL, 100% assay level). %PTMMIV was the average %PTM at the standard injection volume.

### Qualification of NPD

NPD was qualified for the characteristics of limit testing for impurities according to ICH Q2 (R1) guideline, which included specificity and detection limit (Figure 1). To establish NPD specificity, a water injection, blank sample (an aliquot of water prepared side by side with mAb-1 sample following the same sample preparation procedure), mAb-1 RS, mAb-1 RS spiked with 10 pmol of PRTC mixture for each injection, and water injection with 10 pmol PRTC were analyzed by MAM. To determine the limit of detection (LOD) for NPD, different levels of spiked PRTC peptides (0, 0.1, 0.5, 1, 5, 10 pmol) for each LC-MS injection was spiked in a digested mAb-1 sample.

### Proof-of-concept (POC) of MAM NPD for comparability study

Six samples of mAb-1 DS from each of the two different processes (Process 1 and Process 2) were prepared according to the sample preparation procedures.^24^ The samples were run with the same LC-MS conditions and analyzed with NPD in Chromeleon.

## Results

### Qualification of MAM for identity

The specificity of MAM for ID of mAbs were qualified with seven single mAbs and 3 coformulations containing mAb-1. As shown in Table 1, the generic ID method using PARs of N-terminals of both heavy chain and light chain of each molecule to the constant PENNYK peptide of the same sample were able to specifically identify mAb-1 and its co-formulations. To distinguish mAb-1 co-formulations from mAb-1 alone, there were positive IDs for both mAb-1 and the corresponding co-formulated mAb. For mAb-1 single entity, there was only single positive ID for mAb-1 alone. The highest PAR was 0.93 for the positive ID of a co-formulated mAb with extreme disparate concentration difference compared with mAb-1. The results indicated the mAb-1 MAM ID method was indeed specific.

**Table 1.** The specificity of MAM for identify of mAb-1 and its coformulations.

| Sample | Identity (ID) |  |  |  |
| --- | --- | --- | --- | --- |
|  | mAb-1 | mAb-2 | mAb-3 | mAb-4 |
| mAb-1 | Yes | No | No | No |
| mAb-2 | No | Yes | No | No |
| mAb-3 | No | No | Yes | No |
| mAb-4 | No | No | No | Yes |
| mAb-5 | No | No | No | No |
| mAb-6 | No | No | No | No |
| mAb-7 | No | No | No | No |
| mAb-1/mAb-2<br>coformulation | Yes | Yes | No | No |
| mAb-1/mAb-3<br>coformulation | Yes | No | Yes | No |
| mAb-1/mAb-4<br>coformulation | Yes | No | No | Yes |

The Peak Area Ratios (PARs) for Lys-C digested N terminals of both heavy and light chain of mAb-1, -2, -3 and -4 were calculated by comparing to the non-modified PENNYK peptide with the formula as described in the method section. For positive ID of one specific mAb or co-formulations (mAb-1, -2, -3 and -4), the PAR from both heavy and light chain peptides must be $\leq 0.97$ . The highest PAR was 0.93 for the positive ID of a co-formulated mAb with extreme disparate concentration difference compared with mAb-1.

**Table 2.** NPD of MAM for batch comparison.

| Threshold (%) | Process 1 |  |  |  |  |  | Process 2 |  |  |  |  |  |
| --- | --- | --- | --- | --- | --- | --- | --- | --- | --- | --- | --- | --- |
|  | R1 | R2 | R3 | R4 | R5 | R6 | R1 | R2 | R3 | R4 | R5 | R6 |
| 0.01 | 24 | 24 | 24 | 23 | 23 | 22 | 3 | 2 | 2 | 3 | 1 | 0 |
| 0.1 | 3 | 3 | 3 | 3 | 3 | 3 | 0 | 0 | 0 | 0 | 0 | 0 |
| 0.3 | 0 | 0 | 0 | 0 | 0 | 0 | 0 | 0 | 0 | 0 | 0 | 0 |

Six samples of mAb-1 DS were prepared according to the sample preparation procedures from each of the two different processes. The samples were run with the same LC-MS conditions and analyzed with NPD in Chromeleon with different peak detection threshold (0.01, 0.1 and 0.3% of TIC). The NPD for all samples were compared to R6 from Process 2. R=Replicate

### Qualification of MAM for PQA monitoring

The specificity, linearity, range, quantitation limit, accuracy, and precision (repeatability and intermediate precision) of MAM were qualified for PQA monitoring.

#### Specificity

Due to the high resolution (140K) of the MS acquisition setting, the EICs for each peptide did not show peaks at appropriate retention times that were significantly interfering with PTM quantification in water, formulation buffer, and in-process sample buffer injections compared to digested mAb-1 samples (data not shown). There was no major (>0.1%) carry-over for all peptides except for the methionine (M) containing peptides (both the modified and non-modified peptide) in our MAM panel, which are long peptides having 40 amino acids. The M oxidated peptide had about 0.4% carry-over and the corresponding non-modified peptide had about 1.4% carry-over from a blank injection following a digested mAb1 sample. The carryover issue from this long peptide had no impact for all the qualification parameters except assay accuracy (recovery) at its lowest injection level as discussed later. The results indicated that the MAM assay was able to specifically monitor PQAs from mAb-1.

#### Precision

As shown in the supplementary Table 1, the precision was evaluated with repeatability of the method (n=6) and intermediate precision (n=6×6=36). Intermediate precision expresses within-laboratory variations from different days, analysts, columns and MS instruments. The precision results in Figure 2 indicated most PQAs had repeatability within 11% and intermediate precision within 20%. Not surprisingly, there was a clear trend that PQAs having higher percentage of PTM tend to have lower intermediate precision and repeatability (%CV).

**Figure 2:**
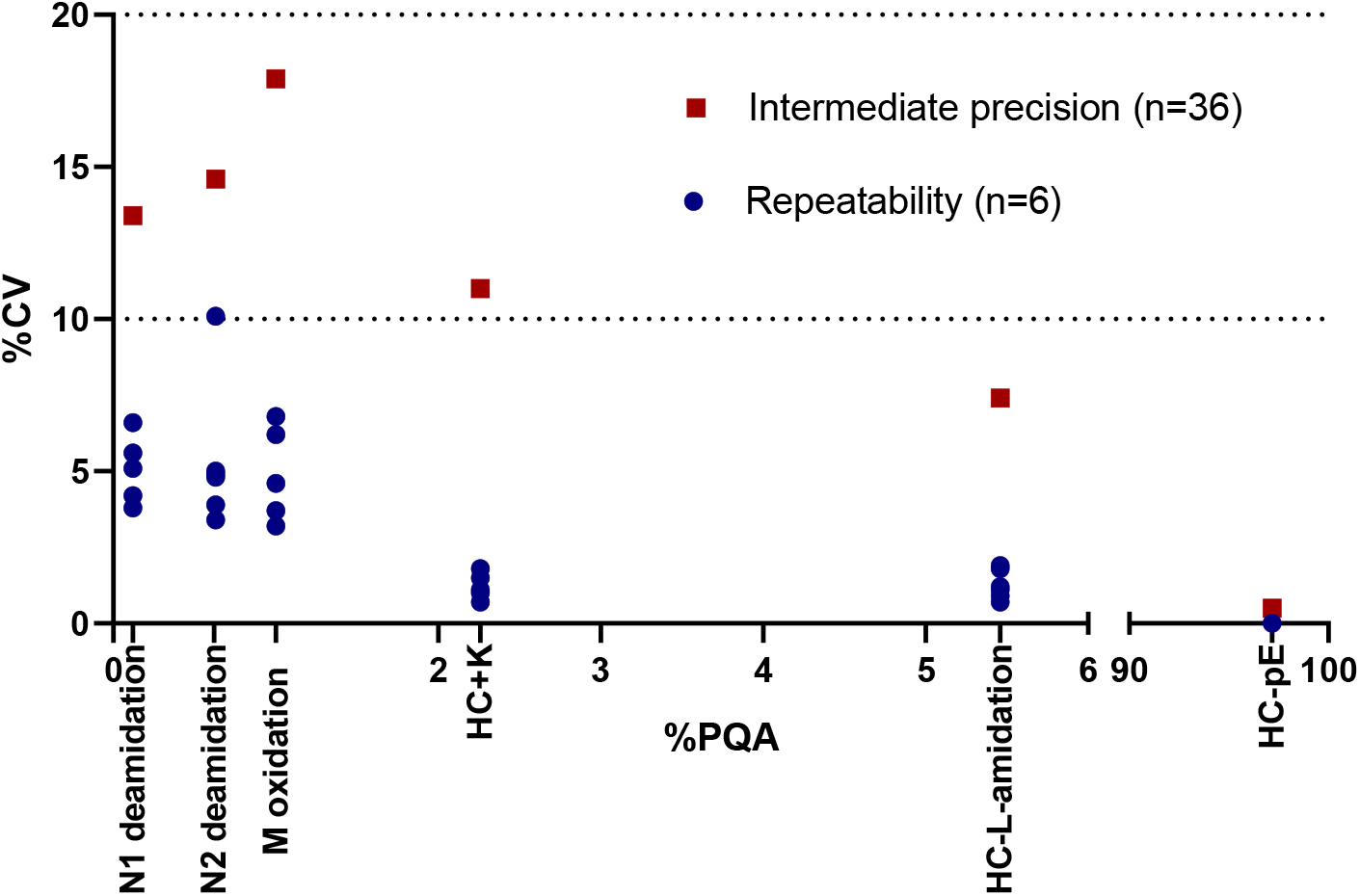
The precision of MAM PQA monitoring. Both repeatability (n=6) and intermediate precision (n=36) were evaluated with two analysts, two columns and two LC-MS instruments using analysis scheme with DOE design as shown in supplementary Table 1. The absolute PTM level of each PQA (%PQA) by averaging all 36 run are shown in the X axis.

#### Linearity

To evaluate the assay linearity, the injection volumes were chosen between 10% and 250% of the target injection volume. As shown in the supplementary Figure 1, all the PQA peptides demonstrated linear relationship between the measured modified or unmodified peak areas against the injection volumes. All the values of goodness-of-fit (R^2^) were above 0.99 across all injection volumes except the N1 deamidated peptide, which only showed R^2^ above 0.99 from 25% to 150% of injection volumes. The narrow linearity range of N1 deamidated peptide was mainly impacted by the autosampler stability of this peptide.^24^ The N1 deamidation level increased across the LC-MS run time of the linearity study.

#### Accuracy

The MAM assay accuracy of each PQA was determined by recovery from the linearity study. As shown in Figure 3, most peptides showed recoveries between 75% to 125%. Because of carry-over issues of the long peptide, the recoveries of M non-modified and modified peptide were over 125% at their 10% assay levels. The recoveries of N1 deamidation peptides were over 125% of recovery at 250% assay level and below 75% of recovery at 10% assay level due to the autosampler stability issue mentioned above.

**Figure 3:**
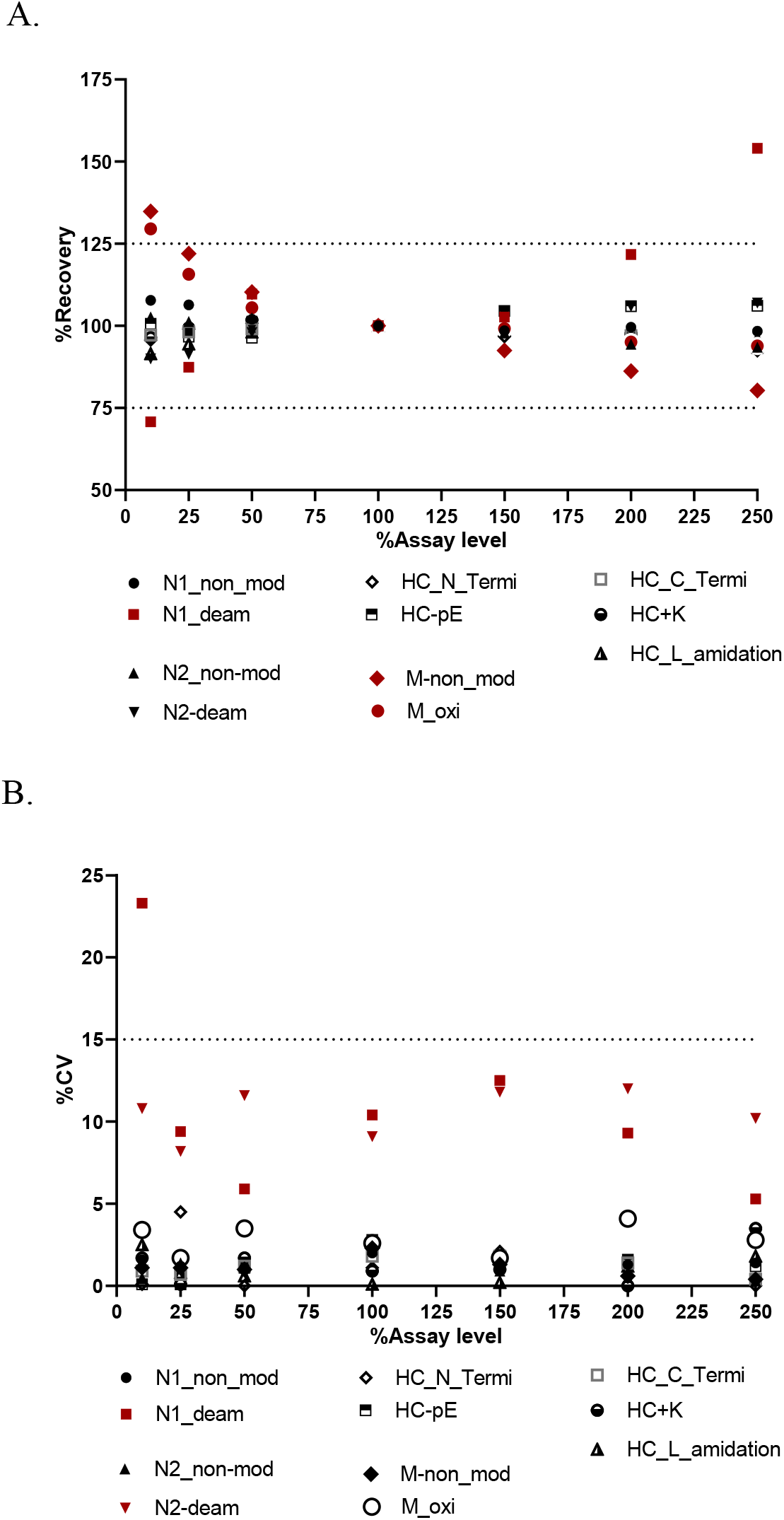
The assay accuracy and precision of PQAs from linearity testing. Assay accuracy (A) were determined by recovery from linearity. The assay precision from the linearity were calculated from the absolute peak areas of each non-modified for modified peptides (n=3).

#### Range

The range of an analytical procedure is the interval between the upper and lower concentration (amounts) of analyte in the sample for which it has been demonstrated that the analytical procedure has a suitable level of linearity, accuracy and precision. The data obtained from linearity study (supplementary Figure 1) of different PQAs were used to determine the assay ranges with suitable accuracy (recovery, Figure 3A) and precision (Figure 3B). Using assay criteria for recovery between 75 to 125% and precision within 15% (%CV), the ranges of most PQAs were from 10% to 250% of assay levels, except the range of M oxidation was from 25% to 250% of assay levels, and N1 deamidation was from 25% to 150% of assay levels.

#### Lower Limit of quantitation (LLOQ)

As shown in the supplementary Table 2, the LLOQ of each PQA was estimated using the formula described in the method section, which used the average %PTM of each PQA observed at 100% assay level multiplied by lowest assay level with acceptable linearity, accuracy and precision. The LLOQ range for all PQAs are from 0.03% to 0.55%.

### Comparison of Convention Methods

In addition to the determination of assay accuracy from the linearity study, the comparison with an independent conventional procedure is another approach to estimate the accuracy of MAM for certain PQAs. The conventional assay, hydrophobic interaction chromatography (HIC)-UPLC, was used to monitor Fab region M oxidation. High levels of M oxidation reduced mAb-1 potency in a forced degradation study (data not shown). The MAM approach of measuring M oxidation using EIC integration adds extra specificity and confidence compared to TIC integration from the traditional assay using UV as a detector. The comparison between HIC and MAM for mAb-1 under different levels of oxidation stress is shown in Figure 4, which demonstrated excellent correlation (R^2^=1.00).

**Figure 4:**
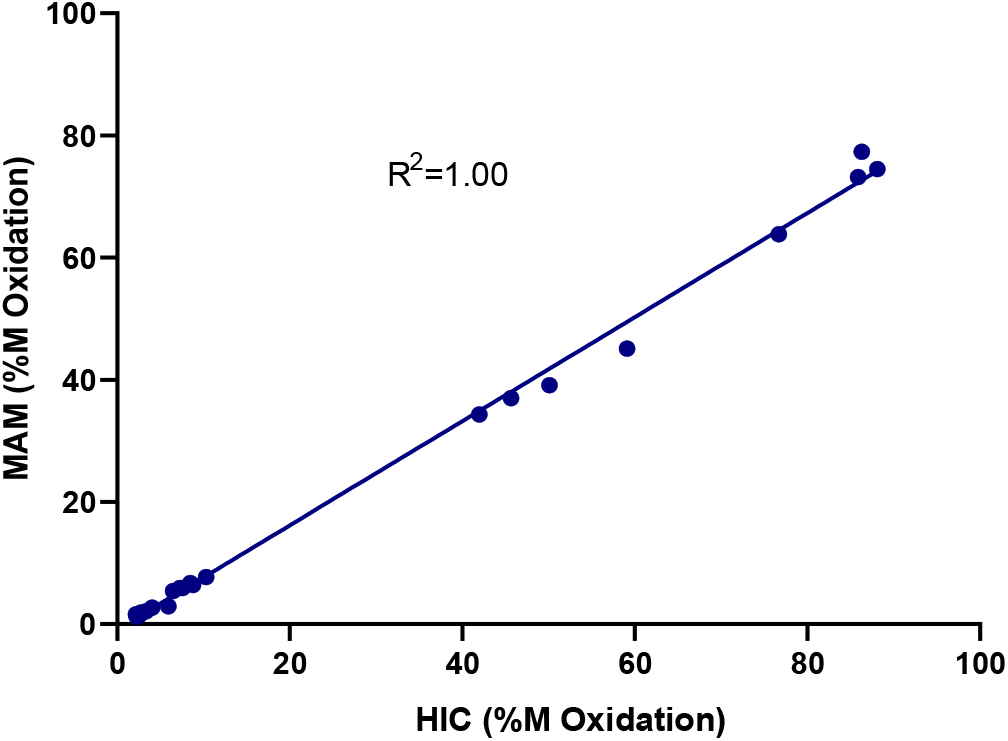
The comparison of M oxidation from MAM with conventional HIC assay. Various stressed mAb-1 samples were tested by both MAM and HIC assays. The %M oxidation were calculated for comparison.

### Qualification of MAM for NPD

The NPD feature of MAM was qualified with its specificity and LOD. To determine the LOD of MAM NPD, different amount of PRTC peptides (0, 0.1, 0.5, 1, 5, 10 pmol) spiked in mAb-1 samples (3.2 μg of mAb-1, about 43 pmol of each mAb heavy chain or light chain) were injected for each LC-MS run. Chromeleon software was used to detect and filter the new peaks from the spiked PRTC peptides (Supplementary Figure 2). As shown in Figure 5, the lower LOD (LLOD) is dependent on the properties of the peptides and parameters used in Chromeleon, especially with peak intensity threshold. There were 3 peptides which were detected at 0.1 pmol (0.2% compared to heavy chain or light chain) spike-in with a 0.01% TIC threshold (Figure 5 and supplementary Table 3). In order to identify all 15 spike-in peptides without false positive peptides, the LOD is 0.5 pmol with 0.01% peak intensity threshold (TIC), and the LOD is 5 pmol with 0.1% and 0.3% of peak intensity threshold (TIC, n=3). When 0.1% TIC was used, 12 out of 15 peptides (80%) were detected for the injection with 1 pmol spike-in (2% compared to heavy chain or LC chain). Not surprisingly, with the lower peak intensity threshold (e.g., 0.01% TIC), more impurity peaks from those peptides or adducts from the 15 spike-in peptides were detected, such as those extra peaks detected from samples with 5 or 10 pmol of spike-in peptides (supplementary Table 3). Most of those impurity peaks were detected in the spike-in peptides when the PRTC peptides were injected alone (Supplementary Figure 2), and were not shown in injections with water. Those impurity peaks may be from the peptide chemical synthesis. Additional sodium adducts of two peptides contributed to the new peaks for both the 1 pmol spike-in at 0.01% peak intensity threshold, and 5 pmol spike-in at 0.1% peak intensity threshold (supplementary Table 3). The number and identity of new peaks detected from the 3 replicates were relatively consistent (Figure 5 and supplementary Table 3).

**Figure 5:**
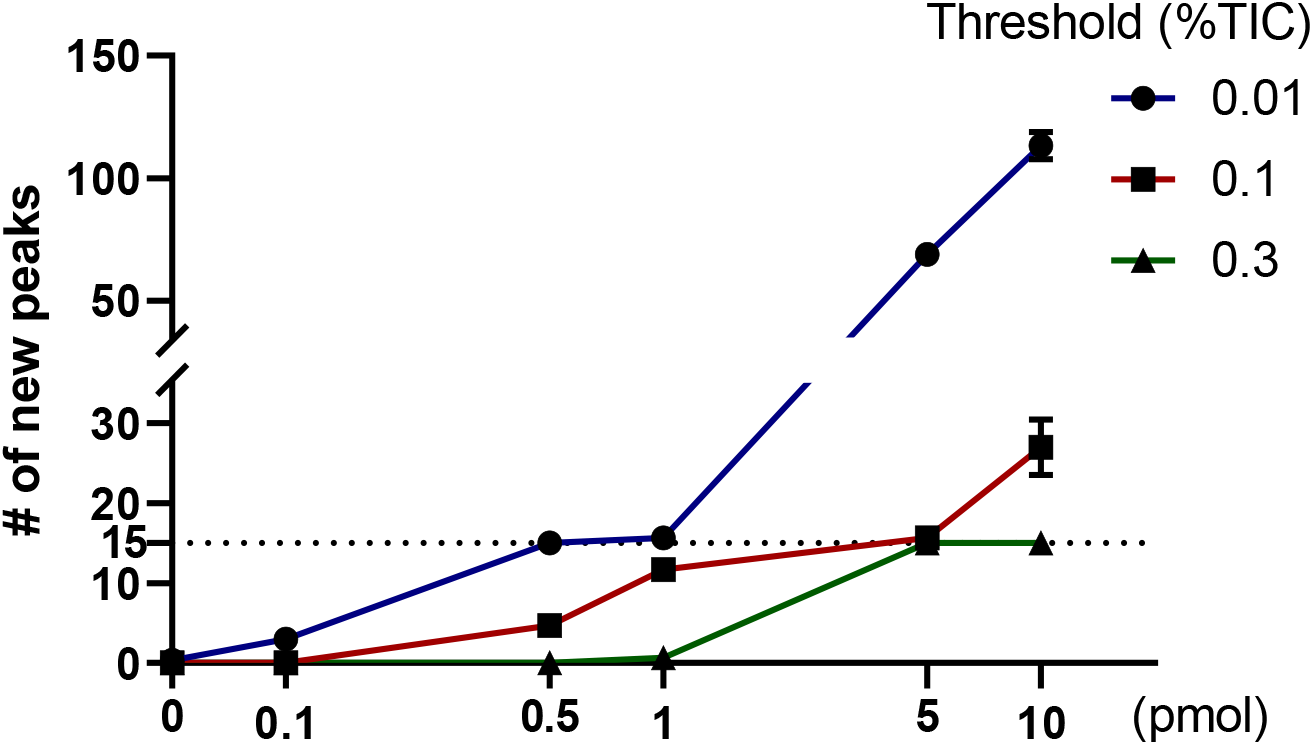
The evaluation of limit of detection of NPD with various amount of PRTC peptides. Various amounts of PRTC peptides from 0 to 10 pmol were spiked in digested mAb-1 samples for each injection (n=3). The detected number of new peaks from Chromeleon were reported. The detailed list of new peaks were shown in supplementary table 3. Data are mean±SD.

### POC of MAM NPD for comparability

The setting of specification for NPD threshold may vary based on sample matrix. To demonstrate the performance of the evaluated NPD parameters in samples without any spike-in peptides, NPD with different thresholds (%TIC) were tested with mAb-1 from two different bioprocesses. As shown in Table 1, more new peaks were detected with the 0.01% TIC threshold in samples from Process 1 compared to those from Process 2, and there were a few (1 to 3) new peaks within different preparations for the same sample from Process 2. With the 0.3% TIC threshold, there were no new peaks within different preparations for samples from both Process 1 and 2. When 0.1% TIC threshold was used, the same 3 new peaks were consistently observed in samples from Process 1 compared to Process 2, and no new peaks within different preparations for the same sample from Process 2. The 3 new peaks were identified as one oxidation-modified and two N-glycosylated peptides, which were consistent with the observations from conventional HIC assay or HILIC-based N-glycan profile (data not shown).

## Discussion

Due to its quantitative nature for specific PQA monitoring and its capability for NPD, MAM is a promising analytical assay for assessing product quality in the product development and manufacturing process. To implement MAM for QC analysis of therapeutic proteins in GMP environment, four aspects need to be carefully considered: (1) risk assessment, (2) method validation, (3) capabilities and specificities of new peak detection feature, and (4) comparison to conventional methods.^6^

### MAM Risk Assessment

Traditional HPLC or CE-based assays provide a general measure of mAb-1 at a global level, but cannot differentiate between species that may overlap using traditional chromatographic approaches.^4–5^ The MAM approach detects more detailed information and higher resolution at the peptide level by measuring site-specific peptide modifications. The NPD feature allows for control of unexpected new modifications, fragments and process-related impurites.^6–7^ The MAM approach, however, may lose information at the intact protein level, and cannot generally tell distribution of modification on the two arms with the same primary sequence using reduced peptide mapping based MAM. Another benefit of MAM is it tests multiple attributes in a single assay, which requires fewer instruments and less footprint. ^6–7^

We envision reduced peptide mapping based MAM can be used in a QC lab as an identity test as well as a purity test, to monitor known CQAs and to detect potential impurities by NPD. A successful transition application of MAM as a primary assay in a GMP lab requires a comprehensive characterization of the therapeutic protein. mAb-1 is a humanized IgG4 antibody. The existing PQA panel covers 2 CQAs (identity and Fab M oxidation) and PQAs related to charge variants which can be captured with conventional assays. The existing identity methods for mAbs include ion exchange chromatography (IEX), potency-based ELISA, and peptide mapping by LC-UV. The use of peptide mapping by LC-MS as an identity method is a logical transition as it can confirm peptides using high resolution accurate mass and retention time (Table 1), which improves specificity over any of those traditional identity methods.

The traditional assay HIC-UPLC was used to monitor a M oxidation on the Fab region as a CQA. High levels of M oxidation reduced mAb-1 potency in a forced degradation study (data not shown). In addition to monitoring the M oxidation in the MAM workbook, the MAM approach can also monitor other oxidation sites. The traditional HIC assay, however, has the ability to monitor M oxidation on different arms of mAb-1 due to increased resolution in HPLC separation by HIC on the intact protein level. However, the HIC assay lacks the ability to localize the oxidation to specific amino acids within the heavy or light chains. The traditional IEX-HPLC assay measures all charge variants in a single chromatogram, which is straightforward for visualization and integration. This conventional assay, however, cannot identify the modifications at a peptide level, which usually requires an additional peak isolation step, followed by MS characterization. The MAM approach has the advantage of detecting specific charge variants on a peptide level, once a full characterization of the molecule has been carried out.

The PQAs associated with the conventional identity, M oxidation, and charge variant assays were selected as proof of concepts for MAM development of mAb-1. Additional PQAs, such as those related to N-glycans, clips, sequence variants, and process residuals can be examined retrospectively with an expanded MAM workbook in the future without additional sample preparation, which is one of the great advantages of MAM.

### MAM method validation

To understand the performance of the improved MAM using Lys-C digestion in non-GMP characterization labs before transferring to QC labs, we performed a fit-for-purpose qualification of the MAM assay for its use of characterization and quality control of therapeutic mAbs. This qualification evaluates the performance of MAM for ID, PQA monitoring, and NPD. As shown in Figure 1, the overall qualification strategy followed the ICH Guideline Validation of Analytical Procedures: Text and Methodology Q2 (R1), which includes the MAM application for ID (identification test), its application for PQA monitoring (quantitative test for impurity), and NPD (limit test for impurity). Thus, only specificity was evaluated for ID, and only specificity and detection limit were qualified for NPD. The assay specificity, precision, linearity, range, accuracy, and quantitation limit were evaluated for PQA monitoring of MAM. There was no quality assurance (QA) approval of the MAM procedures and qualification protocol, and no pre-defined analytical acceptance criteria was set for each assay parameter.

Identity (ID) is one of the most important CQA assessments for biologics release. Traditional ID release methods (e.g., ELISA, IEX, and UV-based peptide mapping) have challenges for co-formulation products, especially for those with extreme ratio differences between two mAbs (e.g., 20:1) or formulations with more than two mAbs. Leveraging the unique N-terminal sequences for both heavy chain and light chain, the PAR approach shown in this study can be considered as a generic or universal approach to specifically identity single mAbs in their coformulations without additional method development (Table 1). This ID approach has not been previously reported in the literature.

The MAM qualification results for PQA monitoring demonstrated its full potential for the intended use. The MAM assay showed great specificity and precision. The assay has excellent linearity across multiple assay levels (injected peptide amount) to have acceptable accuracy and precision. The linearity data indicated the MAM assay can be used to test in-process samples or co-formulation samples, which may have variable protein concentrations for different mAbs, without significant changes to sample preparation, such as increasing protein concentrations to achieve the ideal protein starting amount (100 μg) for each individual mAb in a sample.

The stability of deamidated and oxidated peptides have been a technical challenge for peptide mapping based MS analysis.^4, 19, 25–26^ The baseline level of those PTMs are generally low (< 2%). The issue with autosampler stability has led to higher assay variability compared to other PTMs (Figure 2 and 3). Our optimized digestion protocol using Lys-C digestion for 1 hour without a desalting step was able to control assay precision of those PTMs within acceptable levels (Figure 2). To minimize sample preparation induced methionine oxidation, Carvalho et al demonstrated the benefits of several sample preparation variables for stability of methionine oxidation, which included the decrease of pH during digestion, the addition of methionine at denaturation and after desalting steps.^19^ The low-pH digestion approach together with other digestion parameters (such as different buffer components and addition of ACN) were reported to help control the stability of deamidations. ^25–26^

### Conventional method comparisons

The intent of the MAM assay is to provide orthogonal data sets and ultimately replace conventional assays for ID, charge variants, and oxidation assessment. The extensive product knowledge we gained during mAb-1 development was leveraged for mAb-1 CQA risk assessment when using MAM for in-process, release, and stability testing. To compare back to an orthogonal method for accuracy assessment of MAM for M oxidation, MAM has excellent correlation with the conventional HIC assay (Figure 5). MAM offers additional benefit for site-specific modifications, especially for co-formulated mAbs. The similar approaches will be evaluated and bridged in forced degradation and real-time stability studies with other conventional assays, such as IEX and CE-SDS as reported.^10^

mAb-1 is a IgG4 molecule, thus its N-glycan profile is not a CQA for release testing. N-glycan peptides were not included for qualification. The data collected on the major N-glycan species from the same sample preparation of this MAM assay during process development correlated very well with a conventional method using RapiFluor-MS N-Glycan fluorescence (R^2^>0.99, data not shown). This MAM assay can be easily adapted to quantify N-glycans for other mAbs if N-glycans are deemed as CQAs.

### New Peak Detection

In addition to the targeted monitoring of known PQAs, another important component of MAM is its NPD feature. The purity component of MAM is accomplished by comparing the target molecule to a certified reference standard. The workflow for the purity component of the MAM involves alignment of the chromatograms from the test sample and the reference standard. Detection or absence of peaks in test samples when compared to reference, or significant changes in the ratios of matching peaks is done after peak detection and differential analysis following the peak alignment. One of MAM’s biggest advantages over conventional methods is that MAM can directly characterize an impurity without enrichment. If MS/MS data are acquired, the impurity can potentially be immediately identified. Qualification of NPD is one of the major challenges of MAM, which will need to determine the lower limit of detection with the right thresholds and fold changes that balanced false positives and false negatives.

We evaluated 3 peak intensity thresholds (0.01%, 0.1% and 0.3%), each with a 3-fold or 5-fold changes of peak intensity for the filtering parameters in Chromeleon. The parameter of fold-change had minimal impact on the number of peaks detected by NPD, this may be because we used the spike-in PRTC peptides for this evaluation (Supplementary Figure 2), for which the major changes are obviously larger than 3 or 5 folds. To overcome the limitations of the PRTC mixture, isotopically labeled protein of interest would be a better surrogate for the qualification of NPD. The peak intensity threshold had the largest impact on the number of new peaks detected in our NPD evaluation. Clearly, when assessing the lower limit of detection, more impurities were found as a function of lower peak intensity threshold, regardless of spike-in levels. The number of new peaks and their identities were relatively consistent across replicates (Supplementary Table 3).

To justify the NPD parameters and performance, a proof-of-concept study for process comparability of mAb-1 was tested to correlate with conventional assays for attribute monitoring, such as HIC and HILIC-based N-glycan. The comparability study demonstrated the great potential of our NPD workflow and its parameters for mAb-1 purity testing in QC, which was able to detect the 3 unique peaks leading to the changes of the conventional assays (Table 1). The performance of NPD may benefit from the streamlined Lys-C digestion without an extra desalting step for sample preparation, which introduced less assay variables and resulted in less digested peptides from Lys-C compared to trypsin digestion.^24^ The peptide separation with 0.02% TFA as a mobile-phase additive may also have advantages on consistent peak separation over the well-used mobile-phase additive 0.1% formic acid that other MAM approaches used. ^10–20^

Due to its technical challenges, NPD of MAM was not fully qualified and is rarely evaluated in the literature.^10–20^ Jakes et al applied NPD for problematic HCP detection using a MAM workflow, which was able to successfully detect HCPs down to levels of 100 ppm,^17^ however, the number of other new peaks detected along with peaks from the two HCPs was not disclosed. Recently, MAM Consortium (http://mamconsortium.org/) initiated an NPD round robin (NPDRR) study to survey the current performance metrics of the platform and provide insight for industry members beginning to develop the platform in their laboratories.^14^ Submitted results from 28 participating laboratories (16 participants also provided their corresponding raw data files) identified certain critical elements of NPD, including common sources of variability in the number of new peaks detected, that are critical to the performance of the purity function of MAM. ^14^ Only 54 % of the participants achieved the detection of all designed spike-in peptides without concomitant reporting of false positives, the rest of participants either had false positives or false negatives. This study is crucial to refine the NPD methodology of MAM and accelerate adoption of it into GMP/QC for the management of product life cycles. The limit of detection of the method from each participant cannot be determined as only one spike-in level of PRTC peptides was used, and intra-laboratory variability was also not assessed in sample matrix with spike-in peptides since only one injection of such sample was requested for testing. Both limit of detection and NPD variability were evaluated in our study according to the qualification requirement.

## Conclusions

The improved MAM for mAb-1 was qualified for ID, PQA monitoring, and NPD for its intended use. The qualification results demonstrated the assay’s potential use for mAb-1 characterization and quality control in regulated labs. The developed MAM assay can be considered as a platform assay and easily adapted to other mAbs and co-formulated mAbs. The MAM qualification approach can serve as a reference for other labs trying to implement MAM for characterization and quality control of therapeutic mAbs.

## Acknowledgments

We thank the members of the Analytical R&D Mass Spectrometry, and internal MAM working group for fruitful discussion. The contributions and discussions from Brent Kochert, Xinliu Gao, Larry Wang, and Hillary A. Schuessler are highly appreciated.

## Supplementary Tables and Figures

**Supplementary Table 1.** The analyzing scheme for intermediate precision

| Day | Analyst | Column | Instrument | Lab |
| --- | --- | --- | --- | --- |
| 1 | A1 | C1 | I1 | L1 |
| 2 | A1 | C2 | I2 | L2 |
| 3 | A1 | C1 | I1 | L1 |
| 4 | A2 | C2 | I2 | L2 |
| 5 | A2 | C1 | I1 | L1 |
| 6 | A2 | C2 | I2 | L2 |

**Supplementary Table 2.** The lower LOQ of PQAs monitoring of MAM

| PQA | N1<br>deamidation | N2<br>deamidation | M<br>oxidation | HC+K | HC-L-amidation | HC-apoQ |
| --- | --- | --- | --- | --- | --- | --- |
| LLOQ(%) | 0.03 | 0.06 | 0.25 | 0.23 | 0.55 | 0.28 |

**Supplementary Figure 1.**
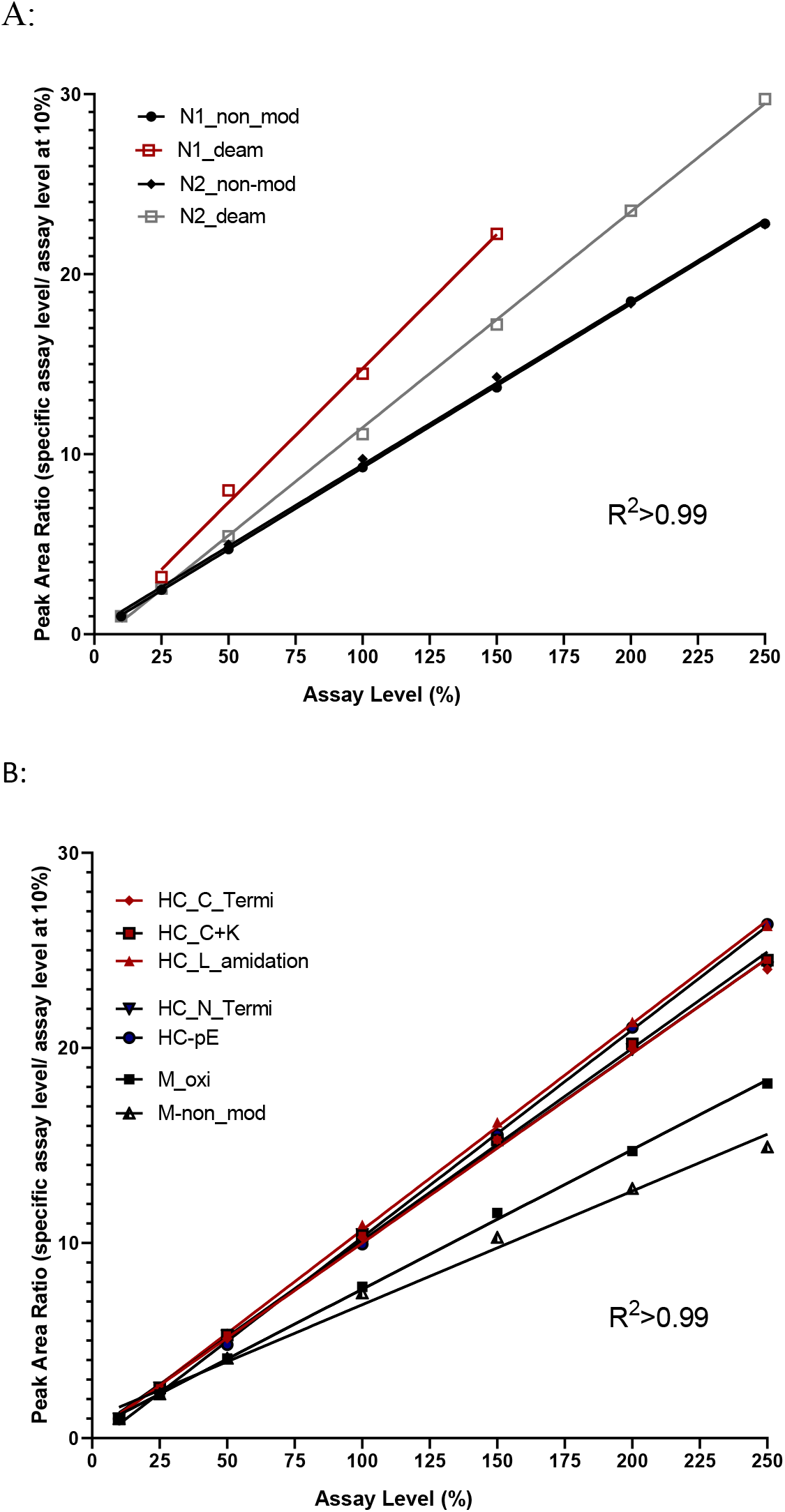
The linearity evaluation of non-modified and modified peptides. The linearity evaluation of non-modified and deamidated peptides from N1 and N2 peptides (A); The linearity evaluation of non-modified and modified peptides from the rest of PQAs (B). Six assay levels from 10% to 250% were tested. The average peak area ratio (n=3) used in the y axis was calculated by peak area of a specific assay level dividing the corresponding lowest assay level (10%). The R^2^ of all fittings are above 0.99 across the labelled assay levels.

**Supplementary Figure 2.**
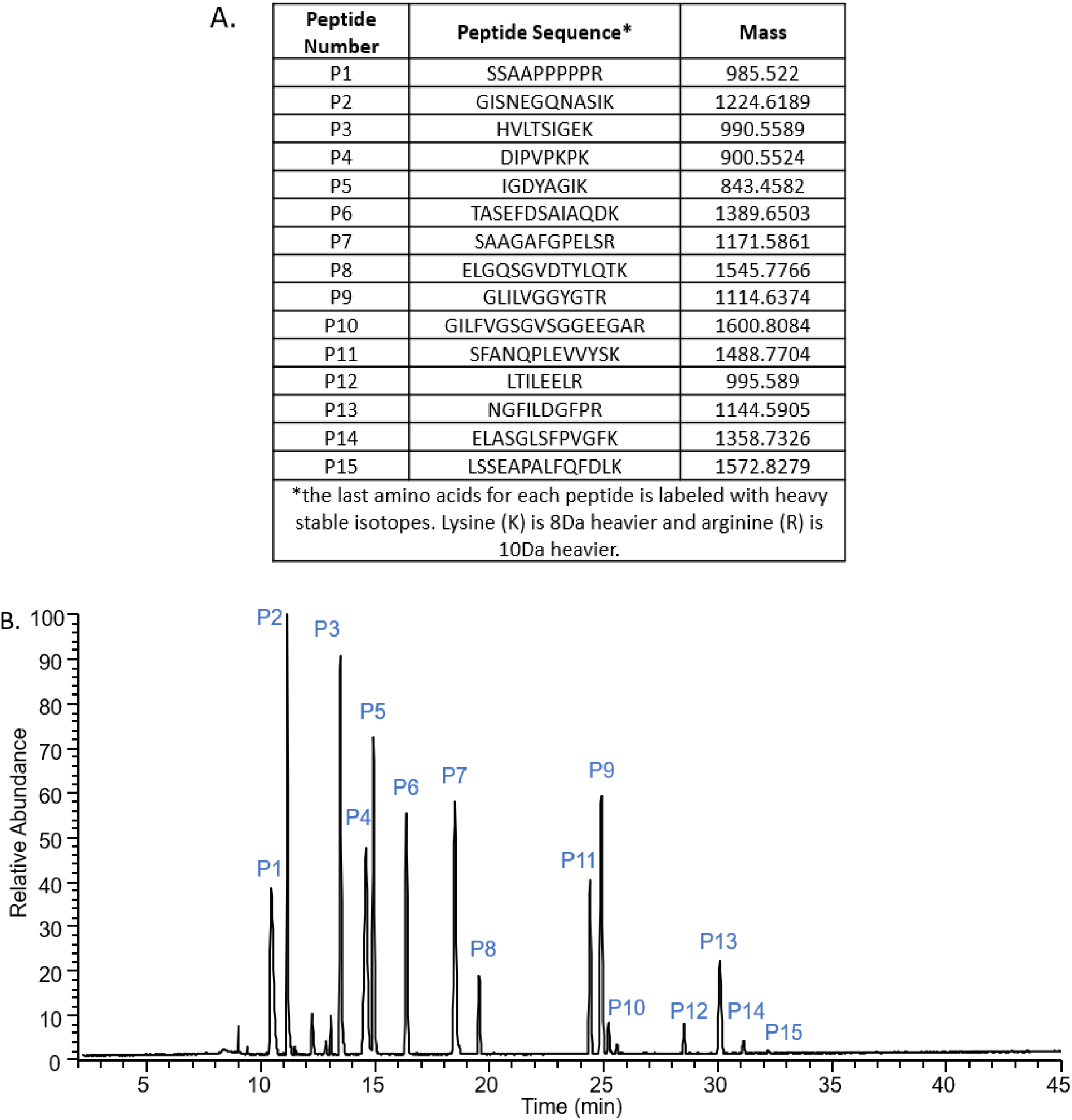
The composition and TIC of the Pierce™ peptide retention time calibration mixture (PRTC) used for NPD. A. The composition of the 15 peptides in the PRTC mixture. B. The TIC of PRTC injection (10 pmol) without any mAb-1 digested matrix run with MAM.

## References

1. Ren, D., Advancing Mass Spectrometry Technology in cGMP Environments. Trends Biotechnol 2020, 38 (10), 1051–1053.

2. Verch, T.; Campa, C.; Chery, C. C.; Frenkel, R.; Graul, T.; Jaya, N.; Nakhle, B.; Springall, J.; Starkey, J.; Wypych, J.; Ranheim, T., Analytical Quality by Design, Life Cycle Management, and Method Control. AAPS J 2022, 24 (1), 34.

3. Rogers, R. S.; Abernathy, M.; Richardson, D. D.; Rouse, J. C.; Sperry, J. B.; Swann, P.; Wypych, J.; Yu, C.; Zang, L.; Deshpande, R., A View on the Importance of “Multi-Attribute Method” for Measuring Purity of Biopharmaceuticals and Improving Overall Control Strategy. AAPS J 2017, 20 (1), 7.

4. Beck, A.; Wagner-Rousset, E.; Ayoub, D.; Van Dorsselaer, A.; Sanglier-Cianferani, S., Characterization of therapeutic antibodies and related products. Anal Chem 2013, 85 (2), 715–36.

5. Beck, A.; Wurch, T.; Bailly, C.; Corvaia, N., Strategies and challenges for the next generation of therapeutic antibodies. Nat Rev Immunol 2010, 10 (5), 345–52.

6. Rogstad, S.; Yan, H.; Wang, X.; Powers, D.; Brorson, K.; Damdinsuren, B.; Lee, S., Multi-Attribute Method for Quality Control of Therapeutic Proteins. Anal Chem 2019, 91 (22), 14170–14177.

7. Rogstad, S.; Faustino, A.; Ruth, A.; Keire, D.; Boyne, M.; Park, J., A Retrospective Evaluation of the Use of Mass Spectrometry in FDA Biologics License Applications. J Am Soc Mass Spectrom 2017, 28 (5), 786–794.

8. Evans, A. R.; Hebert, A. S.; Mulholland, J.; Lewis, M. J.; Hu, P., ID-MAM: A Validated Identity and Multi-Attribute Monitoring Method for Commercial Release and Stability Testing of a Bispecific Antibody. Anal Chem 2021, 93 (26), 9166–9173.

9. Sokolowska, I.; Mo, J.; Rahimi Pirkolachahi, F.; McVean, C.; Meijer, L. A. T.; Switzar, L.; Balog, C.; Lewis, M. J.; Hu, P., Implementation of a High-Resolution Liquid Chromatography-Mass Spectrometry Method in Quality Control Laboratories for Release and Stability Testing of a Commercial Antibody Product. Anal Chem 2020, 92 (3), 2369–2373.

10. Guan, X.; Eris, T.; Zhang, L.; Ren, D.; Ricci, M.; Thiel, T.; Goudar, C., A high-resolution multi-attribute method for product characterization, process characterization, and quality control of therapeutic proteins. Anal Biochem 2022, 114575.

11. Ogata, Y.; Quizon, P. M.; Nightlinger, N. S.; Sitasuwan, P.; Snodgrass, C.; Lee, L. A.; Meyer, J. D.; Rogers, R. S., eAutomated Multi-Attribute Method Sample Preparation using High-Throughput Buffer Exchange Tips. Rapid Commun Mass Spectrom 2021, e9222.

12. Sitasuwan, P.; Powers, T. W.; Medwid, T.; Huang, Y.; Bare, B.; Lee, L. A., Enhancing the multi-attribute method through an automated and high-throughput sample preparation. MAbs 2021, 13 (1), 1978131.

13. Hao, Z.; Moore, B.; Ren, C.; Sadek, M.; Macchi, F.; Yang, L.; Harris, J.; Yee, L.; Liu, E.; Tran, V.; Ninonuevo, M.; Chen, Y.; Yu, C., Multi-attribute method performance profile for quality control of monoclonal antibody therapeutics. J Pharm Biomed Anal 2021, 205, 114330.

14. Mouchahoir, T.; Schiel, J. E.; Rogers, R.; Heckert, A.; Place, B. J.; Ammerman, A.; Li, X.; Robinson, T.; Schmidt, B.; Chumsae, C. M.; Li, X.; Manuilov, A. V.; Yan, B.; Staples, G. O.; Ren, D.; Veach, A. J.; Wang, D.; Yared, W.; Sosic, Z.; Wang, Y.; Zang, L.; Leone, A. M.; Liu, P.; Ludwig, R.; Tao, L.; Wu, W.; Cansizoglu, A.; Hanneman, A.; Adams, G. W.; Perdivara, I.; Walker, H.; Wilson, M.; Brandenburg, A.; DeGraan-Weber, N.; Gotta, S.; Shambaugh, J.; Alvarez, M.; Yu, X. C.; Cao, L.; Shao, C.; Mahan, A.; Nanda, H.; Nields, K.; Nightlinger, N.; Barysz, H. M.; Jahn, M.; Niu, B.; Wang, J.; Leo, G.; Sepe, N.; Liu, Y. H.; Patel, B. A.; Richardson, D.; Wang, Y.; Tizabi, D.; Borisov, O. V.; Lu, Y.; Maynard, E. L.; Gruhler, A.; Haselmann, K. F.; Krogh, T. N.; Sonksen, C. P.; Letarte, S.; Shen, S.; Boggio, K.; Johnson, K.; Ni, W.; Patel, H.; Ripley, D.; Rouse, J. C.; Zhang, Y.; Daniels, C.; Dawdy, A.; Friese, O.; Powers, T. W.; Sperry, J. B.; Woods, J.; Carlson, E.; Sen, K. I.; Skilton, S. J.; Busch, M.; Lund, A.; Stapels, M.; Guo, X.; Heidelberger, S.; Kaluarachchi, H.; McCarthy, S.; Kim, J.; Zhen, J.; Zhou, Y.; Rogstad, S.; Wang, X.; Fang, J.; Chen, W.; Yu, Y. Q.; Hoogerheide, J. G.; Scott, R.; Yuan, H., New Peak Detection Performance Metrics from the MAM Consortium Interlaboratory Study. J Am Soc Mass Spectrom 2021, 32 (4), 913–928.

15. Qian, C.; Niu, B.; Jimenez, R. B.; Wang, J.; Albarghouthi, M., Fully automated peptide mapping multi-attribute method by liquid chromatography-mass spectrometry with robotic liquid handling system. J Pharm Biomed Anal 2021, 198, 113988.

16. Song, Y. E.; Dubois, H.; Hoffmann, M.; S, D. E.; Fromentin, Y.; Wiesner, J.; Pfenninger, A.; Clavier, S.; Pieper, A.; Duhau, L.; Roth, U., Automated mass spectrometry multi-attribute method analyses for process development and characterization of mAbs. J Chromatogr B Analyt Technol Biomed Life Sci 2021, 1166, 122540.

17. Jakes, C.; Millan-Martin, S.; Carillo, S.; Scheffler, K.; Zaborowska, I.; Bones, J., Tracking the Behavior of Monoclonal Antibody Product Quality Attributes Using a Multi-Attribute Method Workflow. J Am Soc Mass Spectrom 2021, 32 (8), 1998–2012.

18. Tajiri-Tsukada, M.; Hashii, N.; Ishii-Watabe, A., Establishment of a highly precise multi-attribute method for the characterization and quality control of therapeutic monoclonal antibodies. Bioengineered 2020, 11 (1), 984–1000.

19. Carvalho, S. B.; Gomes, R. A.; Pfenninger, A.; Fischer, M.; Strotbek, M.; Isidro, I. A.; Tugcu, N.; Gomes-Alves, P., Multi attribute method implementation using a High Resolution Mass Spectrometry platform: From sample preparation to batch analysis. PLoS One 2022, 17 (1), e0262711.

20. Liu, Y.; Zhang, C.; Chen, J.; Fernandez, J.; Vellala, P.; Kulkarni, T. A.; Aguilar, I.; Ritz, D.; Lan, K.; Patel, P.; Liu, A., A Fully Integrated Online Platform For Real Time Monitoring Of Multiple Product Quality Attributes In Biopharmaceutical Processes For Monoclonal Antibody Therapeutics. J Pharm Sci 2022, 111 (2), 358–367.

21. Chauhan, V. M.; Zhang, H.; Dalby, P. A.; Aylott, J. W., Advancements in the co-formulation of biologic therapeutics. J Control Release 2020, 327, 397–405.

22. Kim, J.; Kim, Y. J.; Cao, M.; De Mel, N.; Albarghouthi, M.; Miller, K.; Bee, J. S.; Wang, J.; Wang, X., Analytical characterization of coformulated antibodies as combination therapy. MAbs 2020, 12 (1), 1738691.

23. Patel, A.; Gupta, V.; Hickey, J.; Nightlinger, N. S.; Rogers, R. S.; Siska, C.; Joshi, S. B.; Seaman, M. S.; Volkin, D. B.; Kerwin, B. A., Coformulation of Broadly Neutralizing Antibodies 3BNC117 and PGT121: Analytical Challenges During Preformulation Characterization and Storage Stability Studies. J Pharm Sci 2018, 107 (12), 3032–3046.

24. Li, X.; Rawal, B.; Rivera, S.; Letarte, S.; Richardson, D. D., Improvements on sample preparation and peptide separation for reduced peptide mapping based multi-attribute method analysis of therapeutic monoclonal antibodies using Lys-C digestion. bioRxiv 2022, doi: 10.1101/2022.02.28.482275.

25. Dick, L. W., Jr.; Mahon, D.; Qiu, D.; Cheng, K. C., Peptide mapping of therapeutic monoclonal antibodies: improvements for increased speed and fewer artifacts. J Chromatogr B Analyt Technol Biomed Life Sci 2009, 877 (3), 230–6.

26. Kori, Y.; Patel, R.; Neill, A.; Liu, H., A conventional procedure to reduce Asn deamidation artifacts during trypsin peptide mapping. J Chromatogr B Analyt Technol Biomed Life Sci 2016, 1009-1010, 107–13.

